# Network pharmacology of lycopene and Molecular Docking with Top Hub Proteins

**DOI:** 10.1101/2021.03.04.433249

**Authors:** Nisha Paudel, Umme Hani, Nagendra Prasad Awasthi, Manjunatha Hanumantappa, Rangaswamy Lakshminarayan

## Abstract

**Background:** Lycopene is one of the potent antioxidants in the family of carotenoids that scavenges Reactive Oxygen Species (ROS) singlet oxygen which has been associated with various pathological consequences including atherosclerosis myocardial infarction, and stroke and Sex hormone-induced cancers like breast cancer, endometrial cancer and prostate cancer. As multiple pathways are involved in the manifestation of aforementioned diseases initiated at the behest of ROS, it would be appropriate to understand the likely pathways triggered by the ROS and its eventual control by the action of lycopene through network pharmacology study, a robust paradigm for drug discovery via modulation of multiple targets.

**Results:** 124 proteins were mined from CTD and STITCH databases, which showed some relationship with lycopene, among them strong association was found with TP53, STAT3 and CDK1 proteins. Lycopene showed a strong affinity with these proteins by hydrophobic interactions.

**Conclusion:** The topological analysis of a network created by the lycopene relevant genes showed its role as a potential therapeutic agent in cancer which further requires *in vitro* and *in vivo* studies to confirm these findings.

## 1. Background

Lycopene belongs to the carotenoids family, a class of compounds present in certain fruits and vegetables (1) which is of nutritional interest as its metabolism mediates various signaling effects (2). Lycopene undergoes extensive isomerization, a trans configuration of lycopene is found in fruits and vegetables (3,4) that gives the red hue whereas, it isomerizes in blood plasma and tissues after absorption into cis isomers (2,4). Lycopene is the most abundantly found carotenoid in the human body which is one of the most potent antioxidants that scavenges reactive oxygen species (ROS) singlet oxygen (5). Lycopene is a tetraterpene merged from eight isoprene units that are composed of carbon and hydrogen and it’s an open-polyene chain, without the ionone ring as in β-carotene, gives this molecule with an advantage of a very high antioxidant capacity, which is significantly higher than any other carotenoids, namely, more than twice that of β-carotene and about 10 times higher than that of α-tocopherol (6,7). Particularly, it is effective in the quenching of superoxide anion free radicals, (6). Therefore, its main protective mechanism is in its antioxidant properties, which have been well characterized in vitro (8).

Numerous studies and meta-analysis have been done in the association of lycopene with cardiovascular risks, including atherosclerosis myocardial infarction, and stroke (1,3,9–13) and Sex hormone-induced cancers like breast cancer, endometrial cancer and prostate cancer (14).

Various mechanisms have been postulated by which lycopene alleviates diseases such as prevention of oxidative DNA damage (15) and balancing redox signaling (16), inhibition of androgen receptor signaling (17), inhibition of 5-lipooxygenase (18) and cell cycle regulatory proteins (19). These individual studies verify that instead of a single pathway, multiple mechanisms are involved (20). Oxidative stress is thought to be the main cause for the diseases and lycopene has reactive oxygen stress quenching capacity, as it stands out as a potential balanced redox state of a cell, however, the low bioavailability of lycopene is a major hindrance for its actions (21,22)

Network pharmacology is a new paradigm for drug discovery via modulation of multiple targets rather than a single target, encompassing system biology, network analysis, connectivity, redundancy, and pleiotropy (23). It is used to study compound-proteins/genes-disease pathways, which is capable of describing the complexities of biological systems, drugs, and diseases from a network perspective (24)

In the quest of finding possible approach of the working mechanism of lycopene for its various beneficial properties mentioned above, The present study is carried out, firstly, by extracting all the relevant genes/proteins from databases and then annotating by gene ontology, the network is constructed which showed the interaction in between genes. The network is analyzed and hub proteins are determined and docking of lycopene is done with those significant hub proteins.

## 2. Results

### 2.1 Data curation

Lycopene was found to be associated with 124 curated genes/proteins, Table 1, from the STRING database with a medium confidence level of 0.4 and was reviewed on the UniProt database to create a network.

**Table 1:** List of Lycopene associated Genes

|  |  |  |  |  |
| --- | --- | --- | --- | --- |
| ACHE | C4B | CYP7B1 | JUN | RB1 |
| ADD1 | CASP3 | CYP11A1 | KPNA2 | RDX |
| AGER | CASP8 | DDB2 | LDLR | RPS6KA3 |
| AHR | CASP9 | ESR1 | LIPA | RRM2 |
| AIF1 | CAT | FANCC | LPO | SBP |
| AKT1 | CCND1 | FASLG | MAP2K1 | SCGB1D4 |
| AR | CCNE1 | FOS | MAP2K6 | SELE |
| ATP1A1 | CDK1 | GET1 | MAPK8 | SIRT1 |
| ATP1A2 | CDK4 | GFAP | MGAT2 | SKAP2 |
| ATP1A3 | CDK5 | GJA1 | MICA | SMAD3 |
| ATP1A4 | CDK7 | GJA4 | MLH1 | SMC1A |
| ATP1B1 | CDK9 | GPX4 | MMP2 | SRD5A1 |
| ATP1B2 | CDKN1A | GSR | MSH6 | STAR |
| ATP1B3 | CHGA | HMGB1 | NFE2L2 | STAT3 |
| ATP1B4 | CHGB | HMOX1 | NFKB1 | TFF1 |
| ATP2C1 | CRB1 | HSPA4 | NOS1 | TFP12 |
| ATP2C2 | CTSS | ICAM1 | NQO1 | TGFA |
| BACH1 | CXCL14 | IGF1 | NUP214 | TLR2 |
| BAP1 | CYB5A | IGFR | PAH | TLR4 |
| BAX | CYP1A1 | IL10 | PARP1 | TNF |
| BCL2 | CYP1A2 | IL1B | PCNA | TP53 |
| BCO1 | CYP1B1 | IL2 | PLA2G2A | VCAM1 |
| BDNF | CYP2A4 | IL6 | POMC | VCP |
| BRAF | CYP2E1 | IVL | PTGS2 | ZNF138 |
| C1QA | CYP3A4 | JAK1 | RASSF7 |  |

### 2.2 Network construction and analysis

Based on the STRING database evidence, the network was constructed with lycopene associated proteins to build the protein-protein interaction using Cytoscape v 3.2. The resultant network had 928 nodes and 4390 edges, nodes represent proteins and the edges indicate their relations, Fig 1, represents a summary of simple parameters and ,Table 2, contains a list of genes analyzed by network analyzer based on degree. There significant proteins TP53, STAT3, and CDK1 were selected based on their degree 68, 63, and 45 and average shortest path length 2.9, 2.9, and 3.6 respectively. A radial network, Fig 2, was constructed with three hub proteins.

**Fig 1:**
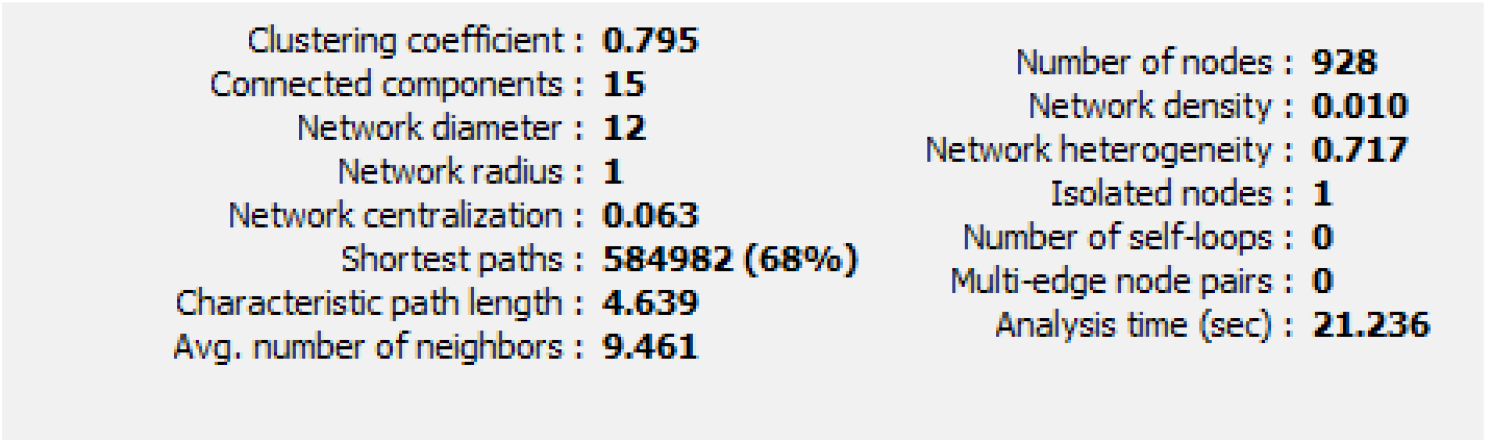
Network Parameters containing the basic details of network

**Table 2:** List of genes based on the degree

| **Name** | **Degree** | **Neighborhood Connectivity** | **Average Shortest Path Length** | **Betweenness Centrality** | **Topological Coefficient** | **Clustering Coefficient** |
| --- | --- | --- | --- | --- | --- | --- |
| TP53 | 68 | 15.6 | 2.9 | 0.19 | 0.05 | 0.11 |
| STAT3 | 63 | 18.63 | 2.9 | 0.15 | 0.06 | 0.15 |
| JUN | 55 | 15.27 | 3 | 0.17 | 0.05 | 0.11 |
| CDK1 | 45 | 13.26 | 3.6 | 0.06 | 0.08 | 0.17 |
| MAPK1 | 45 | 17.08 | 3.1 | 0.07 | 0.07 | 0.19 |
| TNF | 45 | 14.17 | 3.6 | 0.02 | 0.09 | 0.20 |
| AKT1 | 41 | 19.09 | 3.1 | 0.11 | 0.07 | 0.19 |
| MAPK3 | 40 | 15.55 | 3.3 | 0.04 | 0.08 | 0.21 |
| IGF1 | 38 | 13.10 | 3.6 | 0.04 | 0.10 | 0.20 |
| CCND1 | 37 | 20.51 | 3 | 0.04 | 0.07 | 0.27 |

**Fig 2:**
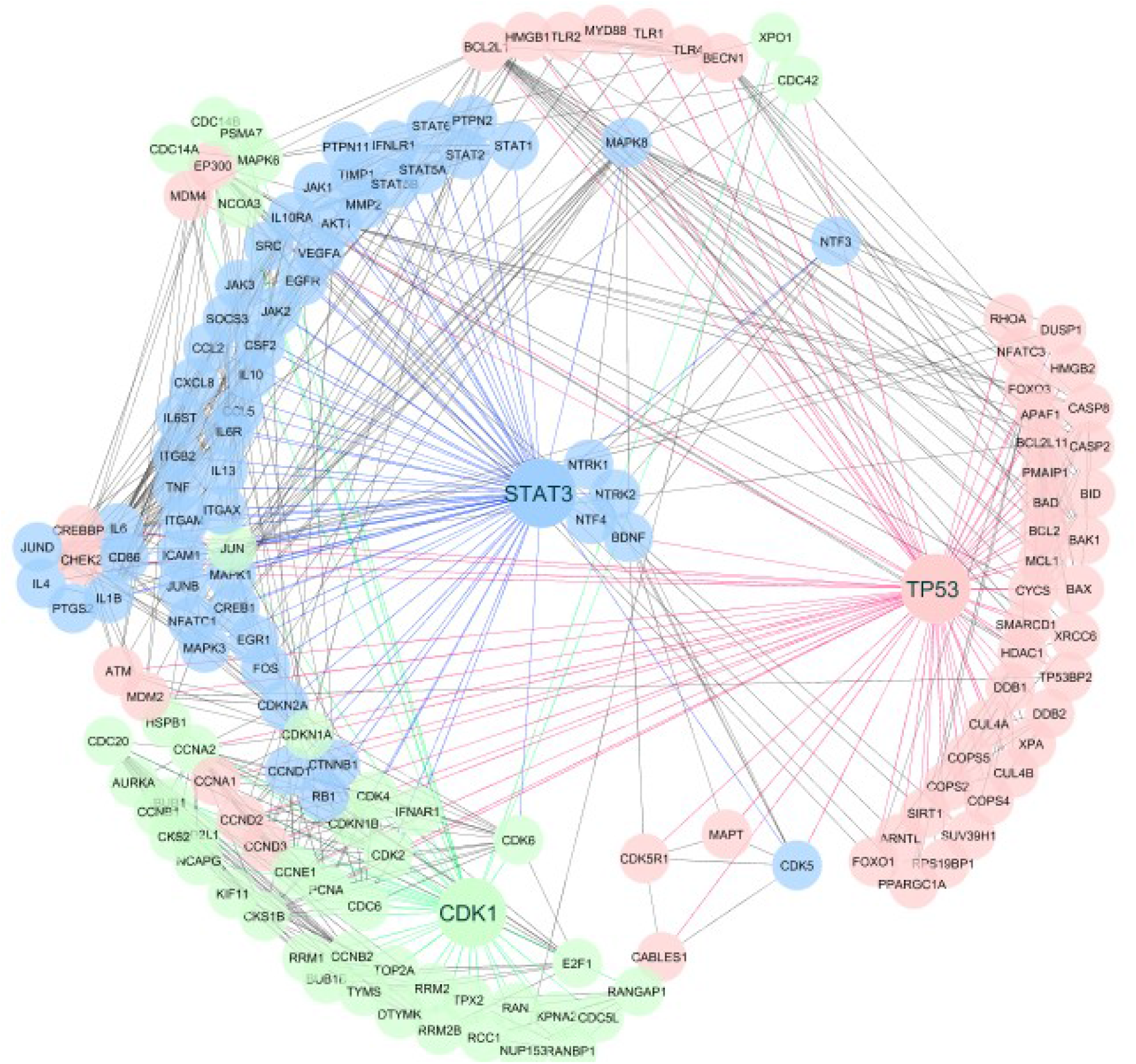
Radial network of genes associated with lycopene constructed by the first neighbor

### 2.3 Gene Ontology analysis

To explore the functional characteristics of lycopene associated genes were subjected to GO analysis and enriched according to the available GO annotations with gene ID and gene accessions in the PANTHER database. In Fig 3 a, molecular function the genes associated with lycopene showed the highest activity in the catalytic activity. In Fig 3 b, the biological process, the associated genes had maximum involvement in the cellular process. In Fig 3 c, cellular component, genes are associated are found in Cell and Cell part. In Fig 3 d protein class, the associated genes had the highest activity in metabolite inter-conversion enzyme and in Fig 3 e, molecular pathway, the associated genes had a major role in CCKR signaling map and apoptosis signaling pathway.

**Fig 3:**
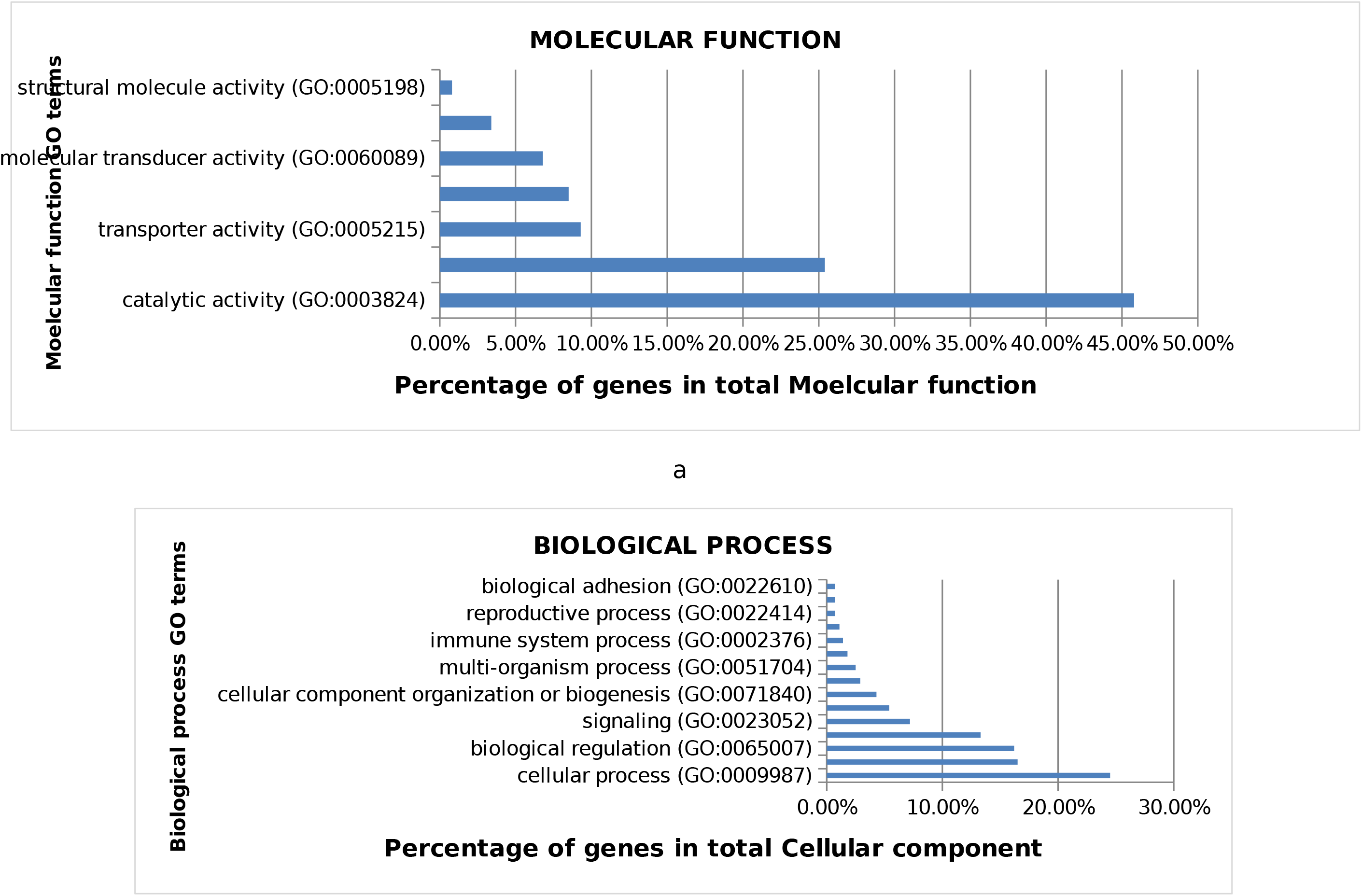
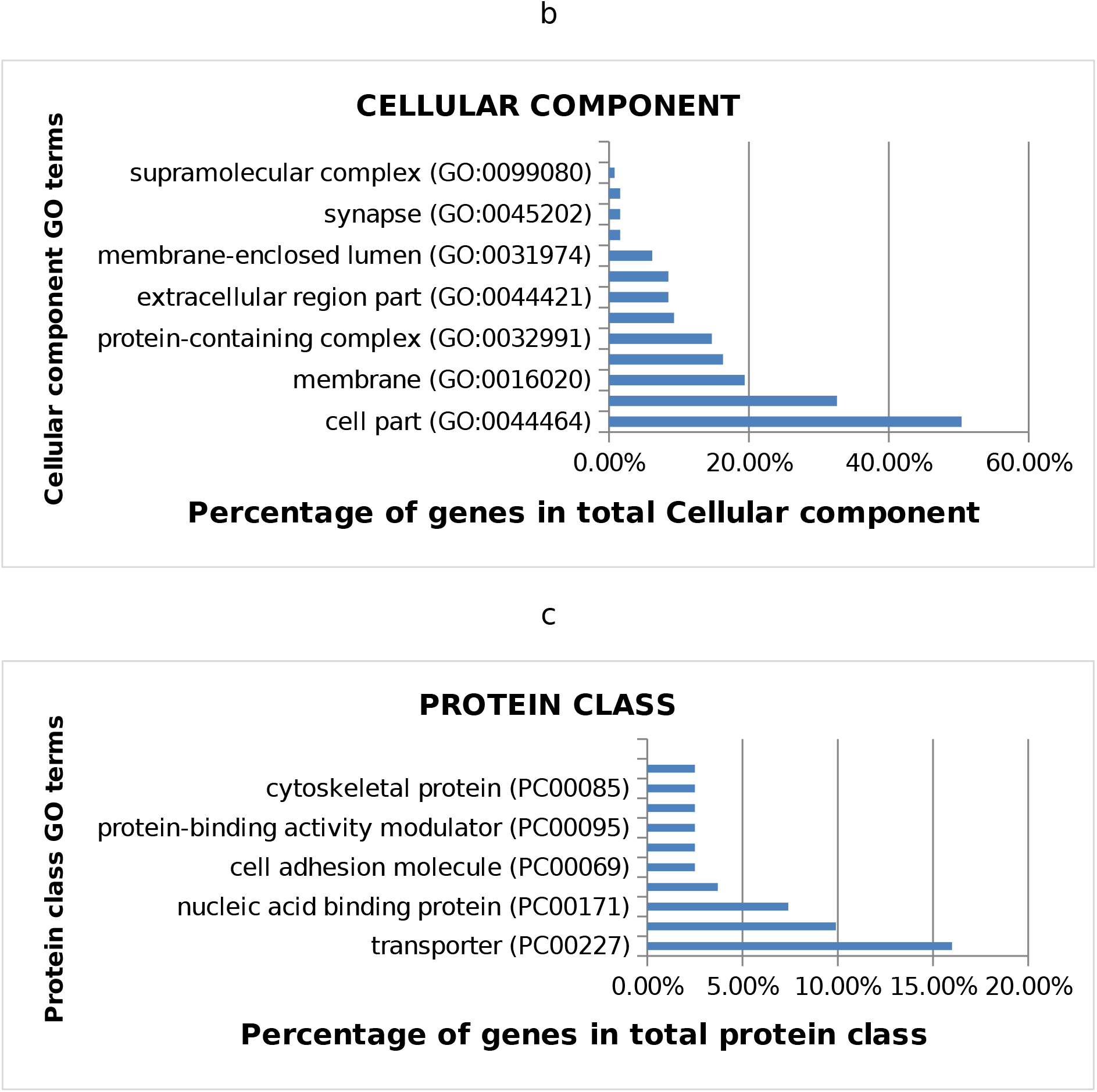
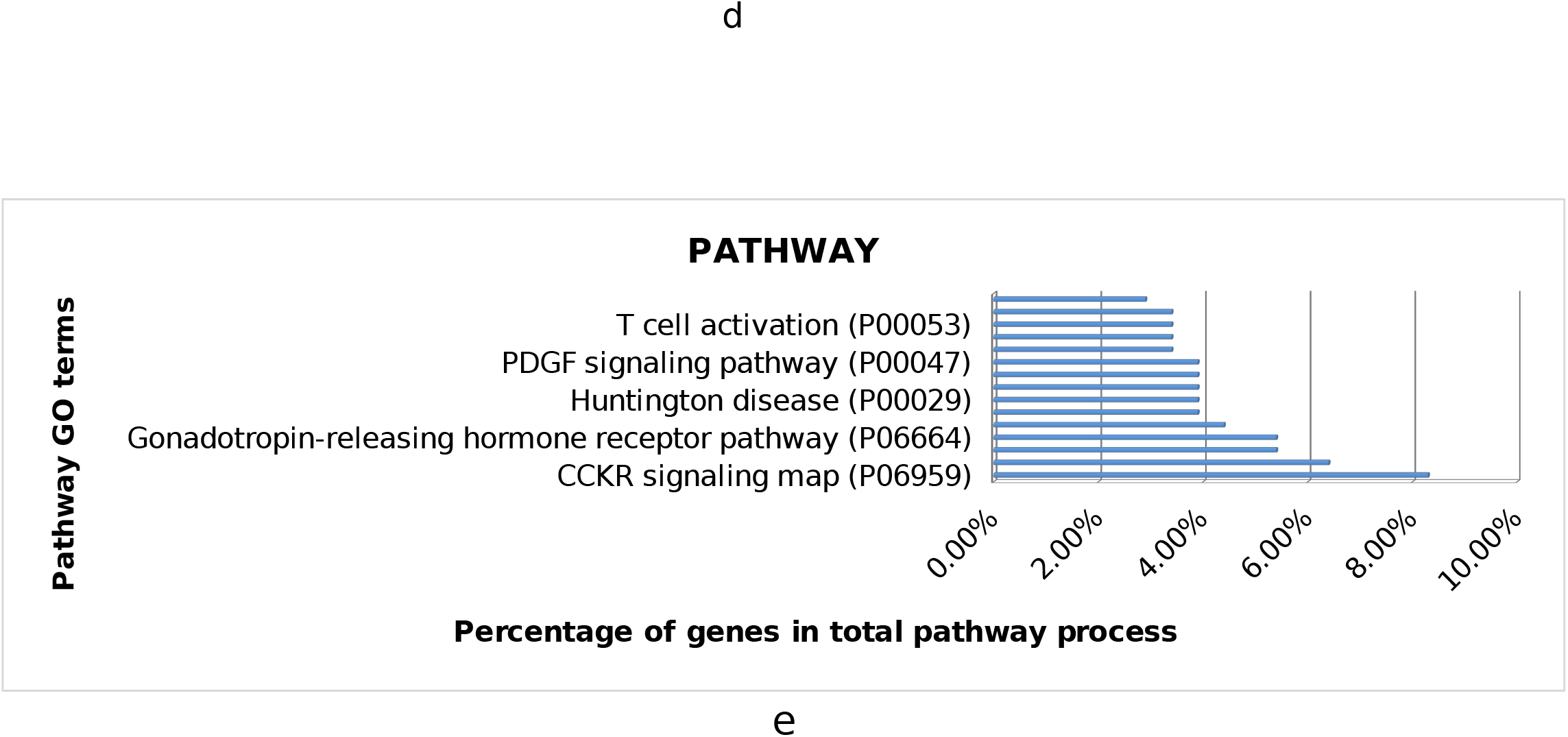
Gene Ontology results a. Molecular function b. Biological Process c. Cellular component d. Protein Class e. Pathway

### 2.4 Molecular docking analysis of lycopene

Top proteins from network analysis were used for docking. Molecular docking analysis of lycopene was performed using Auto Dock Vina Tool and their binding energy was noted. AutoDock Vina results were analyzed based on the interactions between hub proteins and ligand molecule, Table 3. The accuracies of the results were confirmed by considering the lowest binding free energy and hydrophobic binding between the receptor and ligand. The binding energy lycopene with proteins TP53 was − 6.3 kcal/ mol, STAT3 was −6.4kcal/mol and CDK1 was − 5.9 kcal/mol ,Table 3.

**Table 3:** Docking score and hydrophobic bonds of docked complexes

| **Hub proteins** | **Affinity (Kcal/mol)** | **Hydrophobic Interactions** |  |  |  |
| --- | --- | --- | --- | --- | --- |
| TP53 | -6.3 | Lys321<br>Ile332<br>Glu332 | Thr322<br>Phe328<br>Phe338 | Leu330<br>Tyr327<br>Asp352 | Asn345<br>Glu326<br>Asp324 |
| STAT3 | -6.4 | Thr641<br>Gln644<br>Gln635<br>Tyr657<br>Glu625 | Ser614<br>Pro639<br>Val637<br>Tyr640<br>Ser636 | Ile659<br>Trp623<br>Ile 628 | Glu638<br>Asp627<br>Ala662 |
| CDK1 | -5.9 | Arg59<br>Leu58<br>Leu55 | His60<br>Glu57<br>Arg122 | Val124<br>Leu54<br>Glu51 | Arg151<br>Thr47<br>Arg50 |

No, any hydrogen bonds were found in all docking interactions. The hydrophobic interaction profile is shown in Fig 4, Based on the results, it can be inferred that all three proteins can bind with lycopene, which can be further validated by in vitro and in vivo studies.

**Fig: 4.**
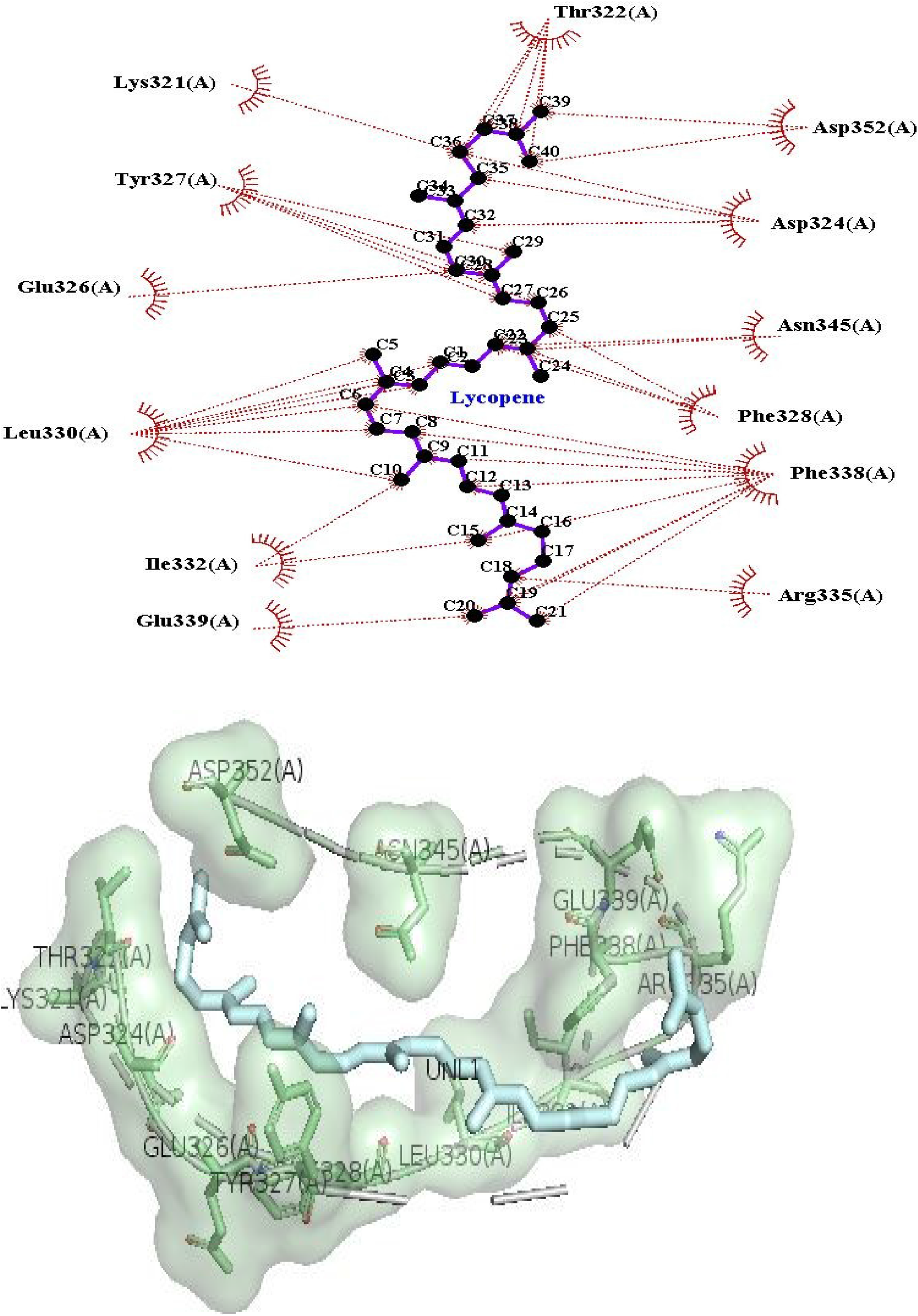
a) Molecular docking TP53-lycopene dock complex ligplot diagram and PyMol presentation respectively

**Fig: 4.**
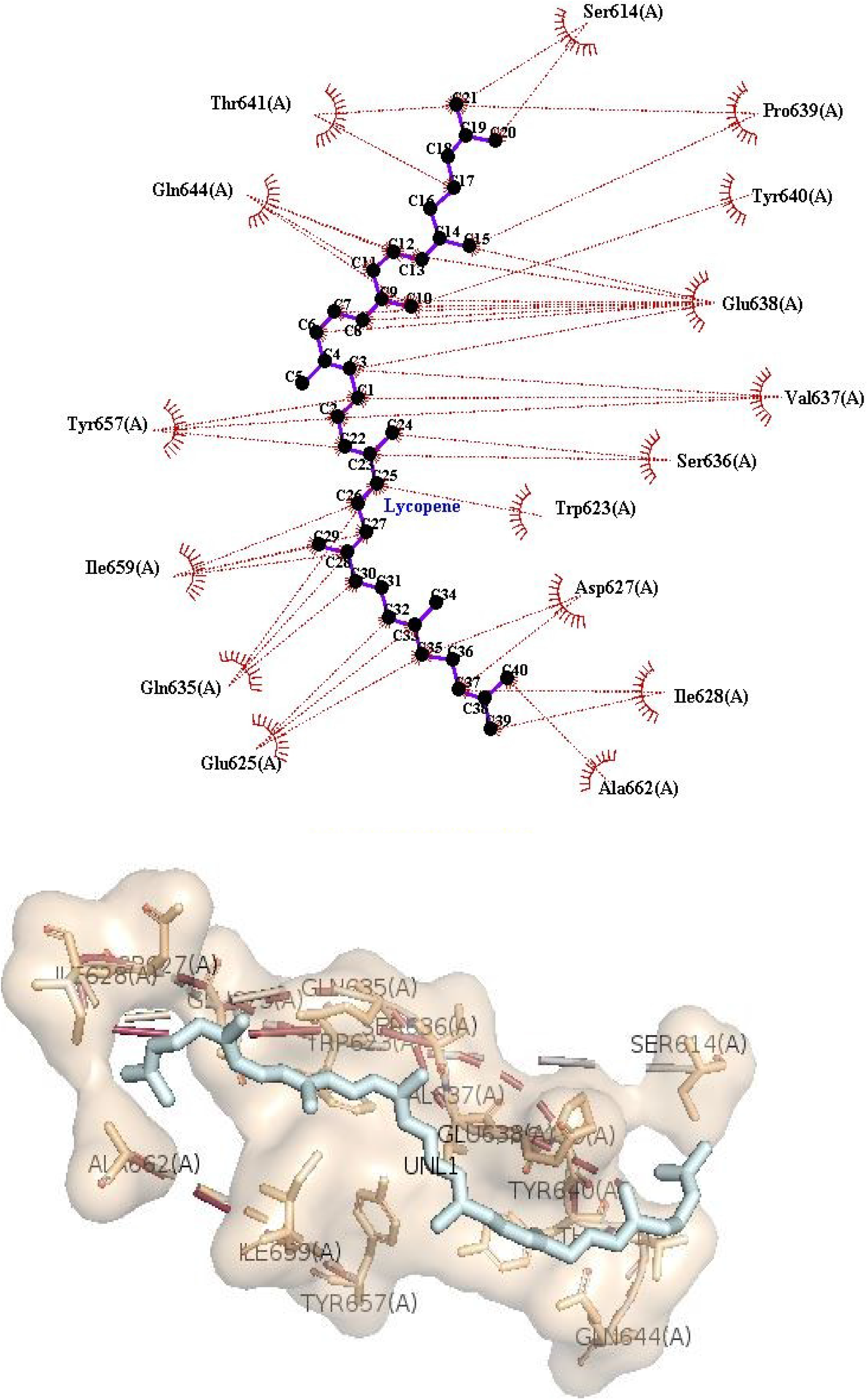
b) Molecular docking STAT3-lycopene dock complex ligplot diagram and PyMol presentation respectively

**Fig: 4.**
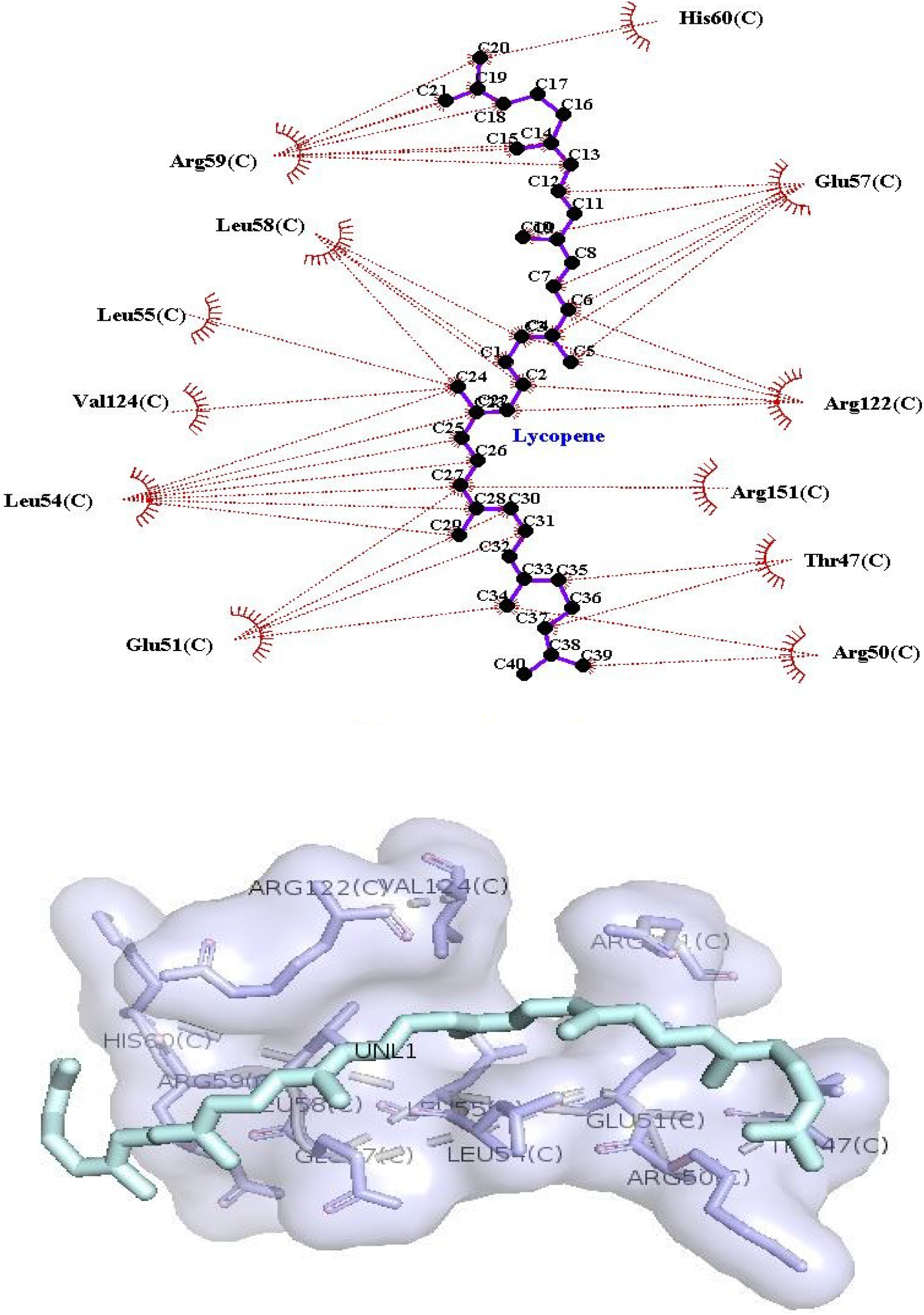
c) Molecular docking CDK1-lycopene dock complex), ligplot diagram, and PyMol presentation respectively

## 3. Discussion

Network pharmacology is a relevantly new emerging technique for drug discovery approach, it’s an effort to address the side effects and toxicity of drug (Hopkins 2007). In the context of lycopene for the first time network pharmacology study has been conducted. Lycopene is a lipophilic, unsaturated carotenoid ranging from yellow to dark red color in various fruits and vegetables.

Numerous studies have been conducted on the beneficial roles of lycopene, such as reducing the risk of some cancers and cardiovascular diseases, however, one of the main obstacles, when considering the biological effects of lycopene, is its very low and variable bioavailability (21,25). The bioavailability depends on various factors including the nature of the double bonds, the geographic sociodemographic factors of an individual, health status, and the food ingestion context (22). These complex factors account for the variability of lycopene concentrations and biological effects in an individual. Furthermore, lycopene distribution throughout the human body is asymmetrical, being mostly found in the liver, adrenals, lungs, prostate, and skin (6). The diseases appear by mediating various signaling effects and the cumulative effect can be studied with the help of network pharmacology.

The study began with data mining from the STITCH and CTD databases and was entered into the STRING database to get relevant genes. Medium confidence (0.400) level was chosen to obtain more number of interactions with affinity. The network was constructed using freeware, Cytoscape, due to the huge number of nodes and edges, which are 928 and 4390 edges, the network did not appear vivid so, a radial layout network was constructed only with the first neighbor nodes of three hub proteins. These three hub proteins were selected based on its topology parameters i.e. degree and average shortest path length; they are TP53, STAT3, and CDK1. All the top proteins obtained had a role in cancer therapeutics. Their structures were retrieved from the RSCB PDB database. 4MZR is a mutated structure of TP53 which no longer protects cells. TP53/P53 is easily mutable protein. So binding of small ligands may stabilize and reactivate the p53 protein activity. As it acts as a tumor suppressor in many tumor types; Involved in cell cycle regulation causes growth arrest or apoptosis depending on the physiological circumstances and cell type (26). 6NJS is STAT3 which is an attractive cancer therapeutic target. Induced degradation of STAT3 results in a strong suppression of its transcription network in leukemia and lymphoma cells degradation of STAT3 protein, therefore, is a promising cancer therapeutic strategy (27). 4YC6 is the structure CDK1 a family of PKs, which plays a key role in the regulation of the cell cycle, they depend on regulatory subunits, cyclin, and their activities are regulated by CDKIPs. In some human cancers, CDKs are overexpressed or CDKIPs are either absent or mutated. Hence, CDKs are appealing therapeutic targets to prevent the unregulated proliferation of cancer cells. As a result, in the last few decades, selective CDK inhibitors have been noticed as effective chemotherapeutic agents (28).

In the simple parameters, obtained after network analysis, the clustering coefficient was found to be 0.795, which signifies at a higher degree, nodes tend to cluster together. The largest distance between the two nodes is 12 which is the diameter of the network whereas the shortest distance between any two nodes is 1 which is the radius of the network. Network centralization is 0.063 which means that the network is more uniformly connected. The nodes have an average shortest path length of 4.639 and at an average number single node has 9.461 neighbors. Network heterogeneity is the tendency of the network to contain the hub nodes which is 0.717.

In the molecular function, the proteins associated with lycopene showed the highest activity in the catalytic activity followed by binding and transporter activity whereas in the biological process the associated proteins had maximum involvement on the cellular process and metabolic process. In the cellular component, these proteins/genes are associated with both cellular and extracellular region which includes plasma membrane and external encapsulating structures. In the molecular pathway, genes showed the highest activity on Protein class CCKR signaling map and Apoptosis signaling pathway, in protein class, the target genes were enriched in metabolite interconversion enzyme.

When the docking was done all the three proteins showed a good affinity with the proteins due to its lipophilic nature with acyclic structure (29), it showed good hydrophobic interactions. In the wet lab studies, using real-time PCR assay, lycopene promoted upregulation of TP53 and Bax transcript expression and also downregulation of Bcl-2 expression in PCa (30). lycopene treatment inhibits the activation of Jak1/stat3 signaling in gastric disease (31). lycopene increases the expression of CDK inhibitors including p21 and p27, as well as the tumor suppressor gene p53, and decreases the expression of Skp2 (32).

It is important to note that the conflicting results about the lycopene also have been published may be related to the wide variety of experimental protocols used to discover any association between lycopene consumption and cardiovascular disease. Furthermore, some have even argued that the sole lycopene is not effective as compared to consuming whole tomato products (25,33).

## 4. Conclusions

The topological analysis of a network created by the lycopene relevant genes showed its role as a therapeutic agent in cancer. Furthermore, the docking results also supported a good affinity binding with TP53, STAT3, and CDK1. Since the bioavailability of lycopene is less and it is converted to a metabolite that may show different affinity with these proteins. Hence, encapsulated lycopene can be used to confirm these in silico findings in vitro and in vivo studies.

## 5. Methods

All the programs were run on a 4.00 GB RAM Intel® Core i3 2.5GHz with a 64-bit Windows 10 Operating System in the HP notebook. The research approach was executed by implementing the following methods:

### 5.1 Data acquisition

For the identification of target genes/proteins associated with lycopene relevant to *Homo sapiens*, data mining was carried from CTD (34) which contains manually curated information about chemical–gene/protein interactions, chemical–disease and gene-disease relationships (35) and STITCH (36) which integrates information about interactions from metabolic pathways, crystal structures, binding experiments and drug-target relationships (37). Genes were curated with a medium confidence score (0.400) to find maximum interactions with affinity. The target genes were screened manually in UniProt (38) and only reviewed one was selected.

Genes obtained from CTD and STITCH databases were enriched in the online tool STRING (39), which aims to provide critical predicted information on protein-protein interactions inferred from genomic resources, experimental shreds of evidence, text mining, co-occurrence and co-expression (40). The data in STRING were weighted and integrated by assigning a medium confidence score of 0.400, which indicates the probability that the interaction is maximum. Such networks can be used for filtering and examining the functional genomics data and for showing an intuitive design for annotating structural, functional and evolutionary aspects of proteins, and also exploring the predicted interaction networks suggests new directions for future experimental research (41).

### 5.2 Gene Ontology analysis

To know the functional association of obtained genes from databases with lycopene, GO enrichment analysis using PANTHER (42) was performed. It is a tool for the classification of genes based on their evolutionary history and functions (43,44). Five aspects of GO determined are; (a) Molecular function where the molecular-level activities performed by gene products, the (b) cellular component represents locations relative to cellular structures in which a gene product performs a function, (c) biological process is for larger processes, or ‘biological programs’ accomplished by multiple molecular activities (45), (d) protein class and (e) pathway.

### 5.3 Network construction

The Protein-Protein Interaction (PPI) network of genes associated with the lycopene molecule was constructed and visualized using open-source Cytoscape v 3.2.2 software. This software provides integrated models for biomolecular interactome networks displaying nodes as proteins and edges as the physical relationship between them (46).

The genes curated were merged using a tool ‘Advanced Network Merge’ and further analyzed by ‘Network Analyser’ to find the topological parameters of the target genes. Top hub proteins were selected and the network was created using the top three hub proteins’ first neighboring proteins. The various parameters of the network were analyzed; like the average shortest path length which is the expected route between the nodes to connect it, betweenness centrality is the number of shortest paths going via a certain node, node degree is the number of edges linked to it, shared neighbor distribution is the maximum number of times, a node shares its neighbor with other nodes, node degree distribution, average clustering coefficient distribution, a topological coefficient is a relative measure for the extent to which a node shares neighbors with other nodes (47). Gene list in descending order with a degree was created along with various parameters of gene-like Neighborhood Connectivity, Average Shortest PathLength, Betweenness Centrality, Topological Coefficient, and Clustering Coefficient. Based on these, three proteins, namely, TP53, STAT3, and CDK1 were selected for docking.

### 5.4 Molecular docking study of the top hub node

The top three hub proteins, namely, TP53, STAT3, and CDK1, identified through network analysis were subjected to molecular docking study. The X-ray crystallographic structure of TP53 (PDB ID: 4mzr, resolution: 2.9 Å), STAT3 (PDB ID: 1WZY, resolution: 2.7 Å) and CDK1 (PDB ID, resolution: 2.6Å) was retrieved from RCSB PDB (48) The energy minimized structures of these compounds were obtained by Swiss Pdb-Viewer v 4.1.0 (49)

The active sites were chosen based on software CASTp (50) and finally confirmed by visualizing them proteins in PyMol v 2.3. Active sites of TP53 are GLY334 ARG337 PHE338 GLU 339 GLU340 PHE 341 ASN 345, STAT3 are SER611 GLU612 SER613 TRP623 PRO629 SER636 VAL637 GLU638 PRO639 TYR640, and CDK1 ILE49 52ILE 53SER 56LYS 69VAL 79LEU. AutoDock Vina program was used over Autodock v 4 to dock ligand molecules with a receptor, as it calculates its grid maps automatically and ranks the results without saving its intermediate results which makes it fast and less space-consuming along with this Vina comes with a new knowledge-based, a statistical scoring function that replaces the semiempirical force field of AutoDock (51). The heteroatoms and water molecules in the protein structure were removed, then hydrogen atoms and, Gasteiger charges were added and torsion degrees of freedom and Solvation parameters were assigned and saved in PDBQT format for further analysis (52).

A target-based docking method was used in this study to prepare the grid maps. The coordinate of the grid box was set at x=30, y=30 and z=30, and the grid center was set at X=−74.881 Y= 23.427 and Z= 6.486 for TP53 , X=13.453 Y= 53.543 and Z= 2.754 for STAT3 and X=−38.074 Y= −14.226 and Z= −12.556 for CDK1. Docking simulation was executed with several modes set to 10 to get more accurate results. The pose with the highest negative Gibbs free energy (ΔG) value was considered as the best conformation and selected for further analysis.

The post-docking analysis was performed using LigPlot v.4.5.3 which displays the protein-ligand interactions mediated by the hydrophobic contacts and hydrogen bonds. In Fig 4, 5 the hydrophobic contacts are represented by an arc with spokes radiating towards the ligand atoms they contact. The contacted atoms are depicted with spokes radiating back. The final result was viewed as the PyMol v.2.3 is a 3D structure visualization tool.

## List of Abbreviations

ROS: Reactive Oxygen Species
CTD: Comparative Toxicogenomics Database
STITCH: Search Tool for Interacting Chemicals
STRING: Search Tool for the Retrieval of Interacting Genes/Proteins
GO: Gene Ontology
PASS: Prediction of Activity Spectra for Substances
PPI: Protein-Protein Interaction
RCSB: Research Collaboratory for Structural Bioinformatics
PDB: Protein Databank
CASTp: Computed Atlas of Surface Topology of Proteins
ΔG: Gibbs Free Energy
Pa: Probabilities of being Active
Pi: Probabilities of being Inactive
CCKR: Cholecystokinin Receptor
TP53: Tumor Inducing Protein 53
STAT3: Signal transducer and Activator of Transcription 3
CDK1: Cyclin-Dependent Kinase 1
CDKIPs: CDK inhibitory proteins
PCR: Polymerase Chain Reaction
PCa: Prostate Cancer

## Acknowledgements

The authors are grateful to Bangalore University Biotechnology Department faculty members.

## Contributions

NP curated data designed the study, NP, NPA and UH cytoscape and docking for multiple times to make it reproducible, NP, HM and RL analysed data, NP, NPA, UH, MH and Rl drafted the manuscript.

## Ethics declarations

### Ethics approval and consent to participate

Not applicable.

### Consent for publication

Not applicable.

## Supplementary information

Supplementary files https://github.com/Nishapaudel/Network-Pharmacolgy

